# On-virion structural dynamics reveal temperature- and receptor-coordinated activation of an alphacoronavirus spike

**DOI:** 10.64898/2026.01.02.697259

**Authors:** Jiaming Liang, Cheng Peng, Yutong Song, Zheyuan Zhang, Jinfang Yu, Xinquan Wang, Sai Li

**Author notes:** These authors contributed equally to this work (Jiaming Liang, Cheng Peng).

## Abstract

Human alphacoronaviruses primarily infect the human upper respiratory tract, yet how physiological temperature and receptor-binding coordinate viral spike (S) protein activation remain unclear. Using cryo-electron tomography and subtomogram averaging, we directly visualized the structural dynamics of S on human coronavirus 229E (HCoV-229E) virions under physiologically relevant temperatures and in the presence of the host receptor human aminopeptidase N (hAPN). We identified two prefusion S2 conformations, compact and loose, that interconvert reversibly by temperature, revealing intrinsic S2 “breathing” on virions. Receptor-binding domain (RBD)-up conformations are rare on apo virions but are promoted at physiological temperature and stabilized upon receptor binding. hAPN can bind bivalently to neighboring S, forming Gemini S-hAPN complexes. Receptor incubation further promotes conversion from prefusion to postfusion S, an effect enhanced at 33. Together, our results demonstrate an optimized temperature for the HCoV-229E viral fitness, establish temperature and receptor binding as synergistic regulators of alphacoronavirus S activation and define an integrated, on-virion model of S conformational transitions during viral entry. These findings provide a structural framework for understanding alphacoronavirus infectivity.

## Introduction

Throughout the last century, human coronaviruses (HCoVs) were primarily associated with mild symptoms of common cold, exemplified by low-pathogenic strains such as HCoV-229E and HCoV-OC43^1,2^. This perception changed markedly following the 2003 outbreak of severe acute respiratory syndrome coronavirus (SARS-CoV), which underscored the capacity of coronaviruses to emerge as high-pathogenic zoonotic agents^3,4^. Subsequent identification of additional HCoVs, including the low-pathogenic HCoV-NL63 and HCoV-HKU1^5,6^, as well as the high-pathogenic Middle East respiratory syndrome coronavirus (MERS-CoV) and severe acute respiratory syndrome coronavirus 2 (SARS-CoV-2)^7,8^, has further catalyzed scientific inquiries into coronavirus-host interactions. These zoonotic spillover events present persistent threats to human health and the global economy. Extensive scientific consensus has been established that certain coronaviruses can cause both chronic and acute organ diseases and exhibit a frequent propensity for cross-species transmission.

Coronaviruses contain four genera: alpha-, beta-, gamma- and deltacoronaviruses. Among the seven known HCoVs, HCoV-229E and HCoV-NL63 belong to alphacoronaviruses, predominantly infect the human upper respiratory tract and circulate widely in human populations^9^. Although these common cold viruses generally cause only mild upper respiratory tract disease in adults, they can lead to severe lower respiratory tract disease in infants, elderly adults, and immunocompromised individuals^10^. However, no approved antiviral drugs or vaccines are currently available for these coronaviruses. Moreover, alphacoronaviruses pose a continued zoonotic risk, as evidenced by the recent identification of emerging alphacoronaviruses in humans presenting with respiratory illness^11–14^. Notably, APN is highly conserved in amino acid sequence across animal species^15^ and serves as a primary receptor of most alphacoronaviruses, including HCoV-229E^16^, porcine coronavirus^17,18^, feline coronavirus and canine coronavirus^19,20^. Together, these observations underscore the need for sustained surveillance and mechanistic studies of alphacoronavirus.

Alphacoronaviruses utilize S protein, a heavily glycosylated homotrimer, to mediate receptor engagement and membrane fusion^21^. The S protein comprises two functional subunits: S1, which contains the N-terminal domain (NTD), the receptor-binding domain (RBD), and the C-terminal domain (CTD) and is primarily responsible for receptor binding; and S2, which drives membrane fusion. Compared with betacoronaviral S, alphacoronaviral S exhibits distinct structural features. In particular, the S1 subunit utilizes an intra-subunit domain packing mode^20,22–27^, in which steric constraints imposed by the NTD limit RBD mobility. The S2 subunit also displays notable conformational plasticity. Previous structural studies of recombinantly expressed HCoV-229E S ectodomains revealed two prefusion S2 conformations: a conventional closed form (S2-compact) and an open form (S2-loose), in which the central helix (CH), heptad repeat 1 (HR1), and fusion peptide (FP) rotate outward relative to the S2-compact conformation. In 2019, an RBD-closed, S2-loose conformation was identified in an S protein stabilized by double-proline (2P) substitutions (T871P and I872P)^25^. In 2021, two additional conformations, including an RBD-closed, S2-compact conformation and an RBD-activated, S2-compact conformation, were resolved from wild-type (WT) S^24^. Structural comparisons revealed the S2-loose conformation exposes the FP^24^, suggesting that the S2 opening may facilitate transition of S from pre- to postfusion state. Recently, a one-RBD-up, S2-loose conformation was determined from an hAPN-bound, 2P-stabilized S, providing insights into receptor-binding geometry^28^. However, structural information for S from other alphacoronaviruses remains limited. For HCoV-NL63, only the RBD-closed S has been reported^27,29,30^, and among animal alphacoronaviruses, an RBD-up state has been resolved only for porcine epidemic diarrhea virus (PEDV) S^31^.

Collectively, current structural insights into alphacoronaviral S remain fragmented. Several fundamental questions therefore remain unresolved: (i) whether alphacoronaviral S can spontaneously adopt RBD-up conformations required to expose receptor-binding sites; (ii) how conformational dynamics of the S1 and S2 subunits are coordinated, and how these states influence receptor-binding competence; and (iii) which factors regulate transitions among the diverse prefusion conformations. Addressing these gaps is essential for establishing a comprehensive mechanistic understanding of alphacoronavirus entry.

To address these questions, we employed cryo-electron tomography (cryo-ET) to directly visualize S on intact HCoV-229E virions under physiologically relevant conditions, including temperature variations and host receptor. subtomogram averaging (STA) analysis revealed temperature-dependent reversible transitions between S2-compact and S2-loose conformations. We further identified two distinct types of S-hAPN complexes on the viral surface. Quantitative comparisons of the ratio of prefusion/postfusion S, and apo/receptor-bound S across experimental conditions revealed the respective contributions of temperature and receptor to shaping the conformational landscape of S. Together, these on-virion structural analyses, involving the membrane context, authentic glycosylation and intermolecular interactions, provide an integrated framework for understanding alphacoronaviral S activation and entry mechanisms.

## Results

### Overview of apo and hAPN-incubated HCoV-229E samples

HCoV-229E infection causes seasonal epidemic, with peak incidence during winter months^9^. During infection and release, HCoV-229E virions experience temperature fluctuations, transitioning from ∼33 in the human upper respiratory tract^32^ to considerably lower environmental temperatures. Given the previous observations that viral replication in nasal cultures is attenuated at 37 relative to 33 ^32^, we first examined whether HCoV-229E propagation in Huh-7 cells was similarly temperature dependent. Virions of the HCoV-229E VR740 strain propagated at 33 exhibited higher titers than those propagated at 37 (Fig. 1a). Consistent with this observation, viral preparations generated at 33 displayed a stronger protein absorption peak and higher overall yields (Extended Data Fig. 1a,b). Therefore, we propagated the viral stocks at 33 for all subsequent experiments.

**Fig. 1.**
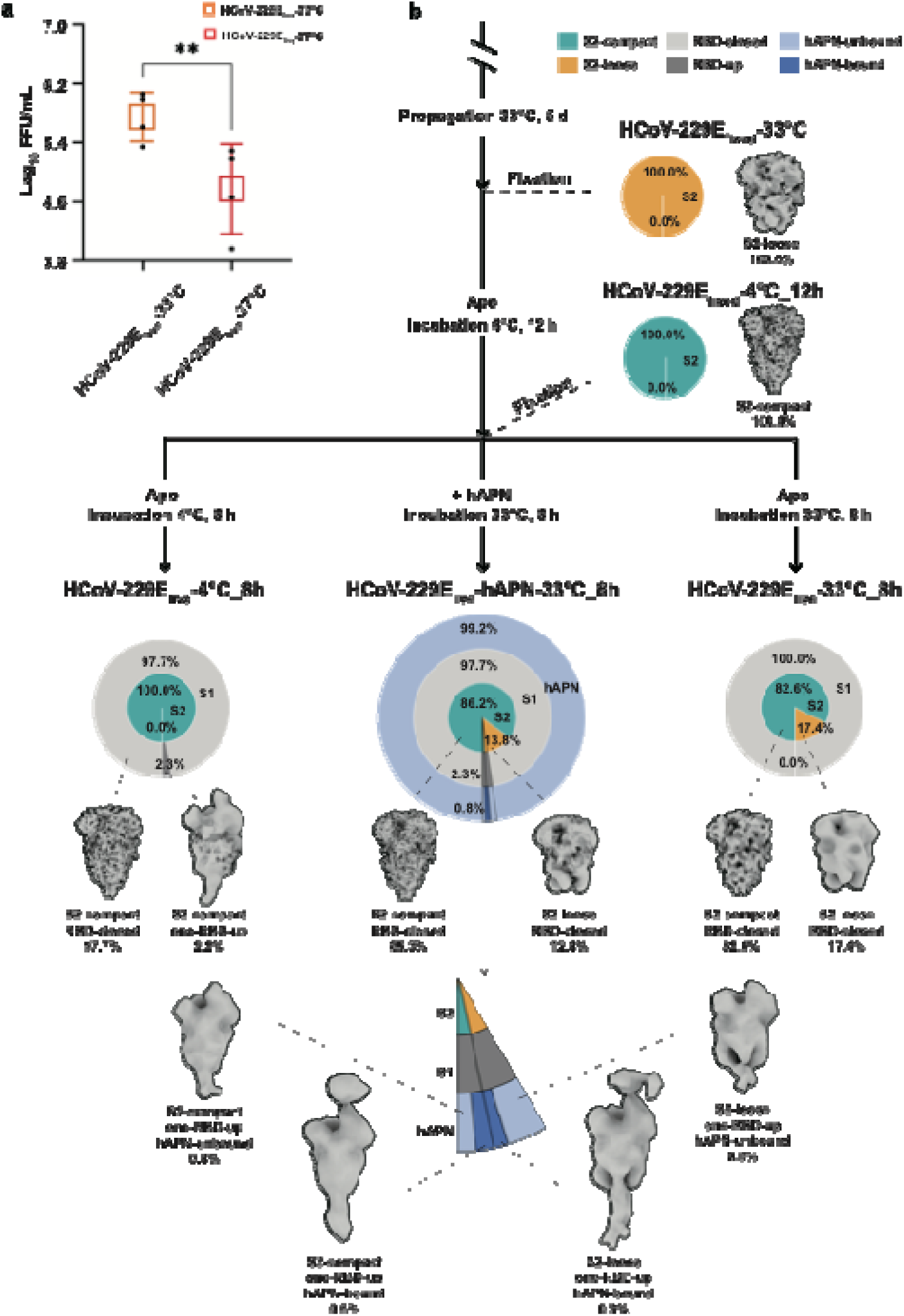
Temperature-dependent infectivity and experimental design for on-virion structural analysis of HCoV-229E spike. **a**, Titers of HCoV-229E virions propagated in Huh-7 cells at 33℃ or 37℃. Virus stocks derived from equal volumes of cell culture supernatants were serially diluted and used to infect Huh-7 cells. Statistical significance was determined by Pair t test (** indicates P value < 0.01). **b**, Schematic overview of sample preparation conditions used to examine temperature- and receptor-dependent conformational dynamics of S on HCoV-229E virions. Virions were subjected to defined temperature treatments (33lJ or 4lJ), fixation protocols, and incubation with or without soluble hAPN. Samples were categorized as apo or hAPN-incubated and as live or fixed, as indicated. The abundances of RBD-closed (light gray), RBD-up (gray), S2-compact (cyan), S2-loose (orange), hAPN-unbound (light steel blue), and hAPN-bound (steel blue) conformations in the viral sample treated with host temperature and receptor (HCoV-229E_live_-hAPN-33lJ_8h) are depicted in a donut chart. Similar analysis for apo viral samples incubated at different temperatures (4lJ and 33lJ) are also shown. The relative maps and their proportions are annotated.

To mimic the physiological and environmental temperatures, virions harvested from cell culture supernatants at 5 days postinfection (dpi) were subjected to defined thermal treatments prior to structural analysis. In one set of conditions, virions were maintained at 33 (HCoV-229E_fixed_-33) or cooled to 4 for 12 h (HCoV-229E_fixed_-4 _12h) before fixation with paraformaldehyde (PFA) and subsequent isolation. To assess temperature-dependent conformational reversibility, a second set of virions was cooled and isolated at 4, followed by incubation at either 33 (HCoV-229E_live_-33 _8h) or 4 (HCoV-229E_live_-4 _8h) for 8 h. To examine receptor-induced effects on S conformation, parallel samples were incubated with the soluble ectodomain of hAPN at 33 for 8 h (HCoV-229E_live_-hAPN-33 _8h). Throughout the manuscript, virions not exposed to hAPN and S not bound to hAPN are referred to as apo virions and apo S, respectively, in contrast to the hAPN-incubated samples. An overview of the experimental design and sample preparation conditions is summarized (Fig. 1b), enabling systematic comparisons of the factors governing on-virion S conformational dynamics.

### The molecular landscape of prefusion S on apo and hAPN-incubated HCoV-229E virions

Cryo-ET and STA of HCoV-229E virions (Extended Data Fig. 2) revealed that HCoV-229E virions displayed an ellipsoidal morphology with envelope dimensions comparable to those reported for other coronaviruses (Extended Data Fig. 3a)^31,33^. Apo and hAPN-incubated samples exhibited comparable virion diameters, prefusion S counts, and the angular distributions of prefusion S tilting relative to the viral membrane (Extended Data Fig. 3a-c). Reconstruction of an exemplary HCoV-229E_live_-hAPN-33 _8h virion exposed to both host temperature showed that the majority of S adopted either an RBD-closed, S2-compact conformation or an RBD-closed, S2-loose conformation (Fig. 2a,b). Apo virions harbored an average of ∼11-14 prefusion S randomly distributed on their surface (Extended Data Fig. 3b), which is fewer than the ∼20 reported for HCoV-NL63^29^ and the ∼26 reported for SARS-CoV-2^33^. Analysis of S orientations demonstrated substantial tilting freedom around the stalk for RBD-closed prefusion S (Extended Data Fig. 3c-e). Across both apo and hAPN-incubated samples, S in the RBD-closed, S2-loose conformation exhibited a modestly broader tilt-angle distribution than those in the RBD-closed, S2-compact conformation (Extended Data Fig. 3c), consistent with increased conformational flexibility in the S2-loose state.

**Fig. 2.**
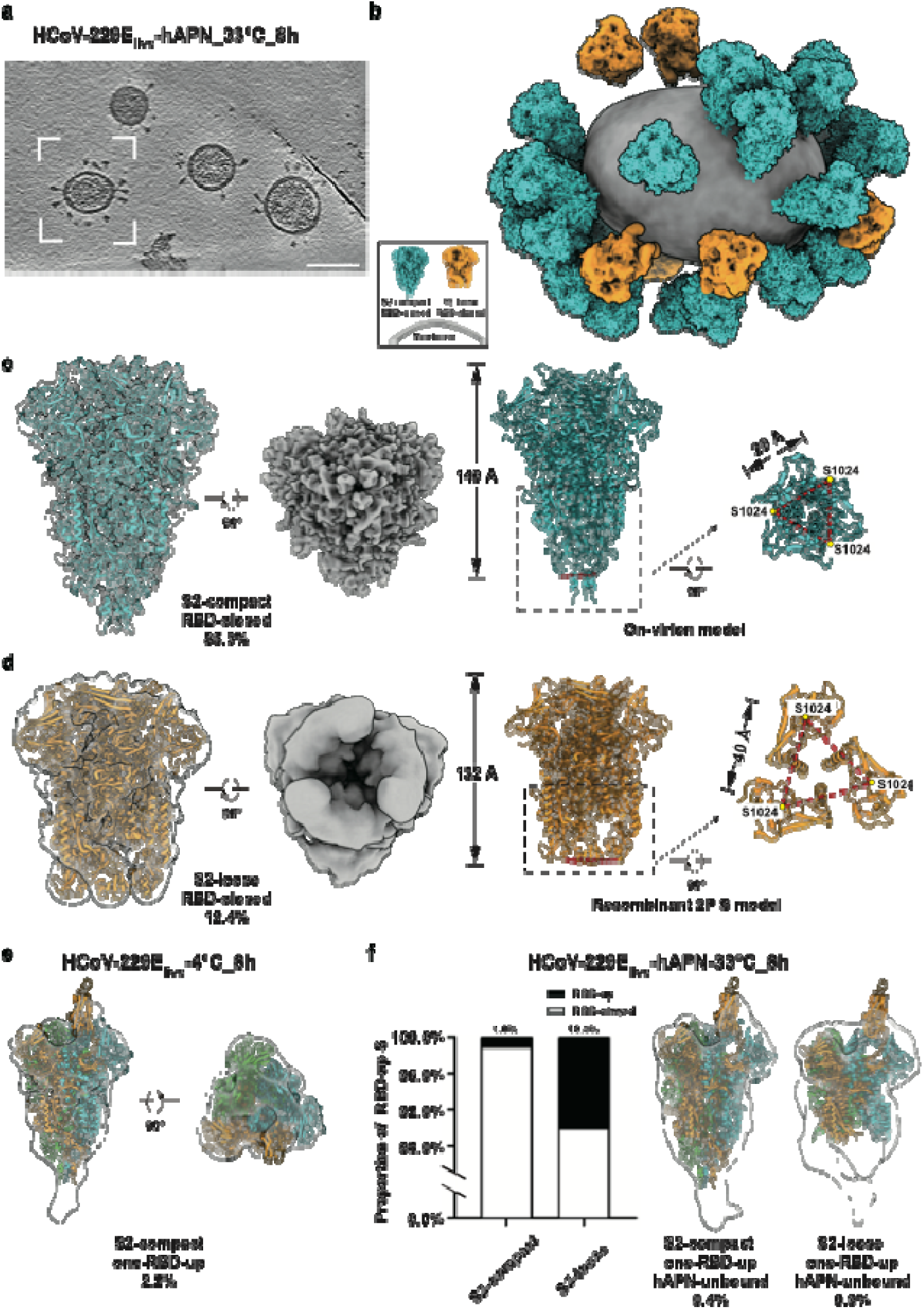
Structural landscape of prefusion S on apo and hAPN-incubated HCoV-229E virions. **a**, An exemplary tomogram slice (14 Å thickness) shows HCoV-229E_live_-hAPN33lJ_8h virions in vitreous ice. Scale bar: 100 nm. **b**, An exemplary virion (boxed in **a**) was reconstructed by projecting the RBD-closed, S2-compact S (cyan), the RBD-closed, S2-loose S (orange), and the lipid envelope (gray) onto their refined coordinates. **c**, STA map and atomic model (cyan) of the RBD-closed, S2-compact S reconstructed from hAPN-unbound S in the HCoV-229E_live_-hAPN-33lJ_8h dataset. The map is shown in side and bottom views. The model dimensions (length excluding stalk, residues 17-1033; width at S2 subunit base, between residues 1024) are shown. **d**, STA map of the RBD-closed, S2-loose S from the same dataset, shown in equivalent orientations and rigidly fitted with the recombinant 2P S model (orange, PDB: 6U7H). The same model dimensions as **c** are shown. **e**, STA map of the one-RBD-up, S2-compact S reconstructed from apo virions (HCoV-229E_live_-4lJ_8h). The map is shown in side and top views. This map was rigidly fitted with the composite model built by superimposing the up-RBD component of recombinant S2-loose S-hAPN model (PDB: 8WDE) onto on-virion RBD-closed, S2-compact model with some modifications. **f**, Left: proportion of RBD-up S among different S2 conformations in the HCoV-229E_live_-hAPN-33lJ_8h sample. States are classified as RBD-closed state (white), RBD-up states (black). Right: STA maps of hAPN-unbound S exhibiting one-RBD-up state in either the S2-compact or S2-loose conformation from HCoV-229E_live_-hAPN-33lJ_8h virions. The S2-compact map (one-RBD-up, hAPN unbound) was rigidly fitted with the composite model built by superimposing the up-RBD component of recombinant S2-loose S-hAPN model (PDB: 8WDE) onto on-virion RBD-closed, S2-compact model with some modifications, while the S2-loose map was rigidly fitted with the S component of recombinant S2-loose S-hAPN model (PDB: 8WDE). The HCoV-229E S-trimer is depicted with its three protomers colored cyan, green, and orange (RBD-up).

### Conformational dynamics of prefusion S on HCoV-229E virions

We next characterized the conformational dynamics of S on intact HCoV-229E virions, revealing intrinsic flexibility within both the S2 and S1 subunits. On the apo-virion sample, S of HCoV-229E_fixed_-33 exclusively adopted an S2-loose conformation, characterized by bifurcation and outward unwinding of the S2 region toward the stalk. In contrast, S of HCoV-229E_fixed_-4 _12h and HCoV-229E_live_-4 _8h exclusively adopted an S2-compact conformation, which is characterized by a canonical closed S2 architecture (Fig. 1b, Extended Data Fig. 3b,d, and Extended Data Table 1). Increasing temperature induced a pronounced shift in the conformational distribution. S of HCoV-229E_live_-33 _8h displayed both S2-compact and S2-loose conformations (Fig. 1b, Extended Data Fig. 3b,d, and Extended Data Table 1). Similarly, following incubation with hAPN at 33, S of HCoV-229E_live_-hAPN-33 _8h again populated both S2 conformations (Fig. 1b, Extended Data Fig. 2, and Extended Data Table 2).

We next examined the conformations of RBD. Since the HCoV-229E_live_-hAPN-33 _8h dataset was substantially larger and the vast majority of S (99.2%) remained unbound to hAPN (Fig. 1b, Extended Data Tables 1,2), we used this dataset to reconstruct apo S structures in both S2-compact and S2-loose conformations (Fig. 2c,d). The S2-compact map, resolved at 4.55 Å resolution, yielded a model highly similar to S2-compact maps derived from apo virions (Fig. 2c, Extended Data Figs. 4 and 5a,b) and to our previously determined on-virion structure from 4 - fixed samples (Extended Data Fig. 5c)^34^.

The highest-resolution features in the S2-compact map were observed in the core ectodomain, whereas the membrane-proximal stalk region displayed lower resolution and lacked discernible density for the transmembrane domain (Extended Data Fig. 4c). Structural alignment with recombinant WT S structures^24^ confirmed close similarity to the RBD-closed state (PDB: 7CYC, RMSD = 1.126 Å) (Extended Data Fig. 6a,b) and greater divergence from the RBD-activated state (PDB: 7CYD, RMSD = 1.377Å). Compared to the recombinant RBD-closed S model, the on-virion model displays an extended helical stalk segment (residues 1034-1047) and increased flexibility on some residues (e.g. residue 312, residue 686) of RBD and S2’, potentially reflecting the flexible nature relevant to further conformational dynamics (Extended Data Fig. 6b). Structural analysis revealed a highly glycosylated map with densities corresponding to twenty-three N-linked glycosylation sites, including four sites (residue 176, 333, 714, 1037) absent from the recombinant WT S structure (PDB: 7CYC) (Extended Data Fig. 6a,c). Rigid fitting of the recombinant 2P-stabilized S model in an S2-loose conformation (PDB: 6U7H) into the corresponding on-virion map produced a satisfactory result (Fig. 2d). Relative to the S2-compact on-virion model, the S2-loose structure is ∼17 Å shorter and ∼11 Å wider, as measured between residues S1024 (Fig. 2c,d), highlighting substantial rearrangements within the prefusion S2 subunit.

We further identified spontaneous RBD-up conformations on HCoV-229E_live_-4 _8h virions. Although rare, 2.3% of prefusion S adopted RBD-up states, predominantly in a one-RBD-up configuration (Fig. 1b). Rigid fitting of a composite model (constructed using the on-virion S model and an RBD-up recombinant S model) revealed an S2-compact, one-RBD-up conformation (Fig. 2e). In contrast, RBD-up conformations were not detected in apo virions exposed to higher temperatures, including HCoV-229E_fixed_-33 and HCoV-229E_live_-33 _8h samples (Fig. 1b and Extended Data Table 1). Following hAPN introduction at 33, however, S in both S2-compact and S2-loose conformations exhibited RBD-up states in the HCoV-229E_live_-hAPN-33 _8h sample (Fig. 1b, Fig. 2f and Extended Data Fig. 2). Notably, RBD-up conformations were modestly enriched in S2-loose S relative to S2-compact S (Fig. 2f). In both S2 conformations, reconstructions of hAPN-unbound S displayed incomplete RBD densities (Fig. 2f), likely reflecting increased conformational heterogeneity or limited particle abundance.

Together, we analyzed the correlation between S1 and S2 conformations with the structures determined from all conditions. These analyses reveal that RBD-up and RBD-closed states coexist on HCoV-229E virions and can occur in both S2-compact and S2-loose conformations. Although RBD-up S represent a minor fraction of the total population, their presence is not restricted to a single S2 conformation, with a modest preference for the S2-loose conformation.

### Temperature drives reversible conformational changes of the S2 subunit

Next, we investigated the factors that promote transitions among the dynamic conformations of S1 and S2 subunits. We conducted a comparative study with three experimental groups to assess the temperature effect on S (Fig. 1b and Fig. 3a). In the first group, in which S conformations were preserved by fixation, we compared the S2 conformations between HCoV-229E_fixed_-33 and HCoV-229E_fixed_-4 _12h samples (Fig. 1b, Fig. 3a and Extended Data Table 1). This analysis revealed that exposure to 4 shifted the S population from the S2-loose to the S2-compact conformation, whereas maintenance at 33 favored the S2-loose conformation.

**Fig. 3.**
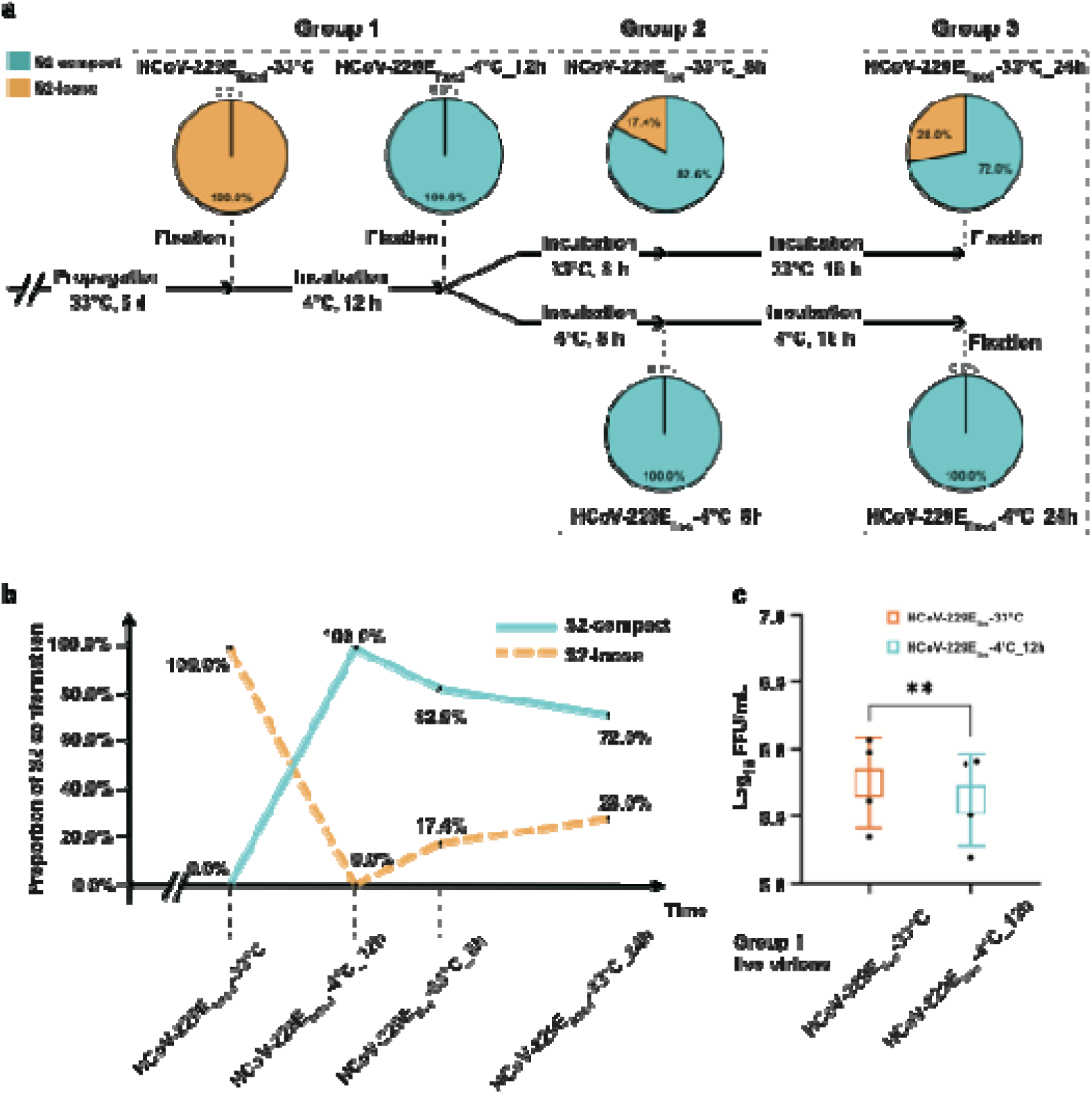
Temperature-driven reversible conformational transitions of the S2 subunit. **a**, Distribution of prefusion S in the S2-compact and S2-loose conformations across HCoV-229E samples subjected to different temperature treatments and fixation protocols, as indicated. Proportion: S2-compact (cyan); S2-loose (yellow). **b**, Chronological alignment of S2 conformational states across sequential temperature treatments. Four samples collected at different time points in **a** are indicated by dashed line on the horizontal axis of the broken line graph. The proportions of S adopting the S2-compact (cyan) and S2-loose (orange) conformations are labeled. **c**, Titers of HCoV-229E virions following temperature treatment. Virions in the left box (HCoV-229E_live_-33lJ) were amplified and maintained at 33lJ, while virions in the right box (HCoV-229E_live_-4lJ_12h) were amplified at 33lJ and then cooled at 4lJ for 12 h until the infection experiment. Statistical significance was determined by paired t test (** indicates P value < 0.01).

To assess whether this temperature-driven transition is reversible, we performed a second comparative group (HCoV-229E_live_-33 _8h and HCoV-229E_live_-4 _8h). STA revealed that warming 4 -incubated virions to 33 for 8 h resulted in reversible conversion of 17.4% of S from the S2-compact to the S2-loose conformation, whereas prolonged incubation at 4 maintained S exclusively in the S2-compact conformation (HCoV-229E_live_-4 _8h) (Figs. 1b and 3a, and Extended Data Table 1). These observations suggest that the low-temperature-driven transition to S2-compact conformation is partially reversible upon exposure to physiological temperature.

In a third group, we examined whether the reversible transition from S2-compact to S2-loose conformations is time dependent. Virions from the second group were further incubated for an additional 16 h at either 33 (HCoV-229E_fixed_-33 _24h) or 4 (HCoV-229E_fixed_-4 _24h) prior to PFA fixation. This operation increased the proportion of S2-loose S from 17.4% to 28.0%, whereas the virions maintained at 4 (HCoV-229E_fixed_-4 _24h) remained exclusively in the S2-compact conformation (Fig. 3a and Extended Data Table 1). These results confirm that extended exposure to physiological temperature progressively promotes conversion from the S2-compact to the S2-loose conformation on HCoV-229E prefusion S.

Across all three experimental conditions maintained at 4 (HCoV-229E_live_-4 _8h, HCoV-229E_fixed_-4 _12h and HCoV-229E_fixed_-4 _24h), S consistently adopted the S2-compact conformation, indicating that neither fixation nor isolation procedures perturbed the 4 -favored state (Fig. 3a). When STA classification results from four sequentially treated samples were aligned along a chronological temperature timeline (HCoV-229E_fixed_-33, HCoV-229E_fixed_-4 _12h, HCoV-229E_live_-4 _8h, HCoV-229E_fixed_-4 _24h), a clear temperature-dependent conformational trajectory emerged (Fig. 3b). Notably, although S2 interconversion is reversible, the transition from S2-loose to S2-compact at 4 occurred more rapidly than the reverse transition from S2-compact to S2-loose at 33.

Previous studies have proposed that the S2-loose conformation represents a primed prefusion state that facilitates progression toward membrane fusion and that receptor engagement preferentially occurs in this conformation^24^. Consistent with this model, we observed that S in the S2-loose conformation exhibited a higher proportion of RBD-up states than those in the S2-compact conformation. Moreover, soluble hAPN was previously shown to suppress HCoV-229E infectivity at physiological temperature but not at 4 ^35^. To directly assess the functional consequences of temperature-dependent S2 conformations, we performed virus infectivity assays using live virions subjected to the same temperature treatments as in the first group (Fig. 3a). Virions maintained at 33 displayed slightly higher titers than those maintained at 4 when briefly adsorbed onto Huh-7 cells (Fig. 3c). This increased infectivity may reflect, at least in part, the higher abundance of RBD-up S and enhanced receptor engagement capacity associated with the S2-loose conformation.

### hAPN receptor induces further conformational changes of S on HCoV-229E virions

Having observed the spontaneous RBD-up states on viral surface (Fig. 2e), we then conducted a comparative study to investigate how receptor binding modulates S conformations (Fig. 4a). hAPN adopts both closed and open states to exert its enzymatic activity^36^, and both states are capable of binding the HCoV-229E S^37^. We first determined the structure of apo hAPN in the closed state at 3.75 Å resolution by single-particle analysis (Fig. 4b), which closely matches the reported crystal structure (PDB:4FYQ, RMSD = 1.283 Å) (Extended Data Fig. 7, Extended Data Tables 4,5). Among the prefusion S on hAPN-incubated virions, the S2-loose conformation contains a higher proportion of hAPN-bound S than the S2-compact conformation (Fig. 4c). The reconstructed S-hAPN maps display stronger up-RBD densities than those observed in hAPN-unbound S (Figs. 2f and 4c), suggesting that receptor binding stabilizes the RBD-up state; and partial hAPN densities, likely due to the low abundance of complex particles. Nevertheless, on-virion S-hAPN maps resemble the corresponding structures determined from recombinant S-hAPN complexes^28^ (Fig. 4c). We tried to extend the incubation of live virions with hAPN at 33 to 24 h (HCoV-229E_live_-hAPN-33 _24h), with a parallel control incubation at 4 for 24 h (HCoV-229E_live_-hAPN-4 _24h), based on previous observations that low temperature stabilizes prefusion states in SARS-CoV-2 S-ACE2 comple^38^. Quantitative analysis indicated that prolonged incubation markedly increased the abundance of S-hAPN complexes, with higher occupancy at 33 than at 4 (Fig. 4a and Extended Data Table 2). These results demonstrate that physiological temperature and extended receptor incubation synergistically enhance S-hAPN binding efficiency.

**Fig. 4.**
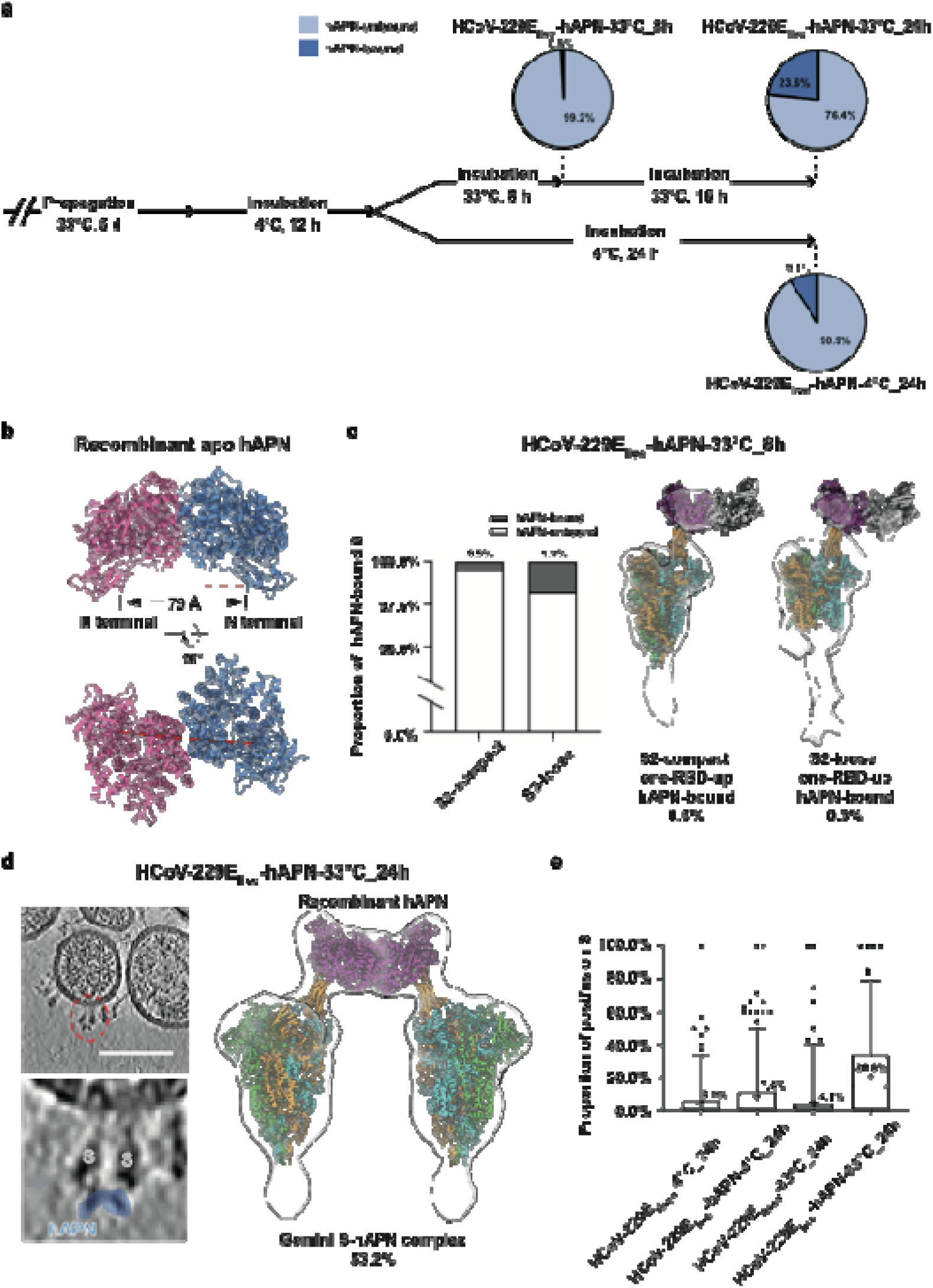
hAPN-induced conformational changes of S on HCoV-229E virions. **a**, Quantification of S-hAPN complex formation under different incubation conditions. Pie charts illustrate their proportion of hAPN-unbound/bound states. Proportion: hAPN-unbound (light steel blue) and hAPN-bound (steel blue). **b**, The atomic structure of soluble hAPN ectodomain dimers. The relative map was determined to 3.75 Å resolution by single-particle analysis. The two protomers are colored pink and blue, respectively. The distance between the N-terminal amino acids of each protomer is indicated by a red dashed line. **c**, Left: proportion of hAPN-bound S across two S2-conformations in the HCoV-229E_live_-hAPN-33lJ_8h sample. States are classified as hAPN-unbound state (white) and hAPN-bound state (dark gray). Right: STA maps of hAPN-bound S exhibiting one-RBD-state in either the S2-compact or S2-loose conformation from HCoV-229E_live_-hAPN-33lJ_8h virions. The S2-compact map (one-RBD-up, hAPN bound) was rigidly fitted with the composite model built by superimposing the up-RBD component of recombinant S2-loose S-hAPN model (PDB: 8WDE) onto the on-virion model with some modifications, while the S2-loose map (one-RBD-up, hAPN bound) was rigidly fitted with recombinant S2-loose S-hAPN model (PDB: 8WDE). The HCoV-229E S-trimer models are depicted with its three protomers colored cyan, green, and orange (RBD-up). The hAPN monomers are colored magenta (bound to S) and white (unbound). **d**, Left: an exemplary tomogram slice (14 Å thickness) of the 24 h incubated HCoV-229E_live_-hAPN-33lJ_24h shows Gemini S-hAPN (red circle), where two S are bridged by an hAPN dimer (light blue). Scale bar: 100 nm. Right: STA map of Gemini S-hAPN complex resolved from HCoV-229E_live_-hAPN-33lJ_24h virions. The map was rigidly fitted with the composite model built by superimposing the recombinant apo hAPN ectodomain model and the up-RBD of recombinant S2-loose S-hAPN model (PDB: 8WDE) onto on-virion RBD-closed, S2-compact model with some modifications. This model is colored as in **c**. **e**, Quantification of prefusion and postfusion S on apo and hAPN-incubated HCoV-229E virions under different temperature conditions. Boxplots display the 5% outliers, minimums, first quartiles, medians, third quartiles, and maximums of the data. The average proportions for each condition are annotated.

At lower contour thresholds, inspection of the averaged prefusion S map from the HCoV-229E_live_-hAPN-33 _24h dataset revealed extended densities bridging adjacent S on virions, forming a second S-hAPN complex, as observed in the tomogram slice (Fig. 4d). Subsequent classification identified two distinct configurations: a Solo S-hAPN complex (46.8% of prefusion S particles), in which a single hAPN molecule binds to one S-trimer; and a Gemini S-hAPN complex (53.2% of prefusion S particles), in which a single hAPN molecule binds bivalently to two neighboring S (Fig. 4d, Extended Data Fig. 8a and Extended Data Table 2). These observations provide direct on-virion evidence for the previously proposed Gemini S-hAPN structure derived from recombinant systems^28^. For the on-virion S-hAPN complex, we performed rigid fitting of the composite model (constructed using the on-virion S2-compact S model and closed apo hAPN model) (Fig. 4d), as well as the recombinant 2P-stabilized S-hAPN model (PDB: 8WDE) (Extended Data Fig. 8b), in which the S adopts an S2-loose conformation and hAPN exhibits an open state. Despite classification and refinement efforts, we were unable to resolve high-resolution maps that definitively distinguished S2 conformations or hAPN states within the complexes, indicating that hAPN-bound S exhibit pronounced flexibility.

Receptor incubation also markedly promoted the appearance of postfusion S on virions (Fig. 4e and Extended Data Fig. 9a). A low-resolution postfusion S map was rigidly fitted with the full-length postfusion SARS-CoV-2 S structure (Extended Data Fig. 9b). Analysis of apo-virion groups (HCoV-229E_fixed_-4 _24h and HCoV-229E_fixed_-33 _24h) indicated that temperature alone exerted only a minor effect on the abundance of postfusion S on virions (Fig. 4e). In contrast, in the presence of hAPN, the proportion of postfusion S increased by ∼90% at 4 and by ∼4-fold at 33 (Fig. 4e). These quantitative results demonstrate that physiological temperature and receptor binding act synergistically to drive the pre-to-postfusion transition of S.

## Discussion

In this study, we utilized cryo-ET and STA to characterize the structural dynamics of S on HCoV-229E virions. We demonstrate that prefusion S simultaneously populate both S2-compact and S2-loose conformations on the viral surface and that these states interconvert reversibly in a temperature-dependent manner. In addition, we identify rare but detectable RBD-up states on apo virions and resolve the structures of RBD-closed and RBD-up S, S-hAPN complexes, and postfusion S under defined thermal and receptor-binding conditions.

Previous studies based on recombinant S ectodomain reported either two S2-compact conformations for the WT^24^ or a single S2-loose conformation for the 2P mutant^25^, but did not capture the coexistence of both S2 conformations within the same S genotype. Our on-virion analyses reveal that S2-compact and S2-loose conformations coexist on HCoV-229E virions after a temperature shift from 4 to 33. To date, RBD-up states in the absence of receptor binding had not been observed in two endemic human alphacoronaviruses, HCoV-229E^24,25^ and HCoV-NL63^27,29,30^. The only reported one-RBD-up structure of HCoV-229E S was derived from a 2P-stabilized recombinant S in complex with hAPN, in which the S adopted an S2-loose conformation^28^. Therefore, how HCoV-229E S engage with receptors remained unclear. Our data reveal that spontaneous RBD-up S can arise on apo virions at 4, although such states were not detectable at 33. While we cannot exclude the possibility that increased conformational flexibility at physiological temperature limits detection of RBD-up S, our cryo-ET analyses clearly demonstrate that, upon receptor incubation at 33, S in both S2-compact and S2-loose conformations can adopt RBD-up states and bind hAPN. This observation contrasts with conclusions drawn from recombinant systems^28^ and underscores the importance of studying S dynamics in their native virion context.

Our results also provide direct on-virion evidence for the existence of Gemini S-hAPN complexes, in which a single hAPN molecule binds bivalently to two neighboring S. Several factors may contribute to these differences observed between on-virion and recombinant systems^24,25,28^. First, our study employed full-length WT S, whereas some recombinant studies relied on 2P mutants designed to constrain prefusion conformations. Second, most recombinant analyses were conducted at 4, whereas our experiments incorporated physiologically relevant temperature (33). Third, the spatial constraint, tilting and diffusion of S on viral membrane^39–41^ likely facilitate hAPN bivalent binding on virions, whereas recombinant systems depend on Brownian motion and concentration gradients in solution for encountering.

Integrating our structural observations across experimental conditions, we propose a temperature- and receptor-dependent model for S activation in HCoV-229E (Fig. 5). At low temperature (4), prefusion S predominantly adopt an RBD-closed, S2-compact conformation stabilized by intra- and inter-subunit interactions of S2 subunits. Upon exposure to physiological temperature (33), this equilibrium gradually shifts toward the S2-loose conformation as thermal energy lowers the energetic barrier for S2 opening. In both S2 conformations, S can spontaneously sample RBD-up states, enabling receptor binding and formation of Solo or Gemini S-hAPN complexes, analogous to SARS-CoV-2 S interactions with ACE2 dimers^38^ and with IgGs^41^. Receptor binding further promotes S1 dissociation and S2 structural rearrangement, ultimately driving membrane fusion and conversion to the postfusion state. The emergence of Gemini S-hAPN complexes may facilitate coordinated conformational transitions among neighboring S, thereby accelerating fusion kinetics. Together, our findings demonstrate that a productive transition from pre- to postfusion state requires coordinated action of physiological temperature, receptor engagement, and sufficient incubation time, reducing the likelihood of premature postfusion transition.

**Fig. 5.**
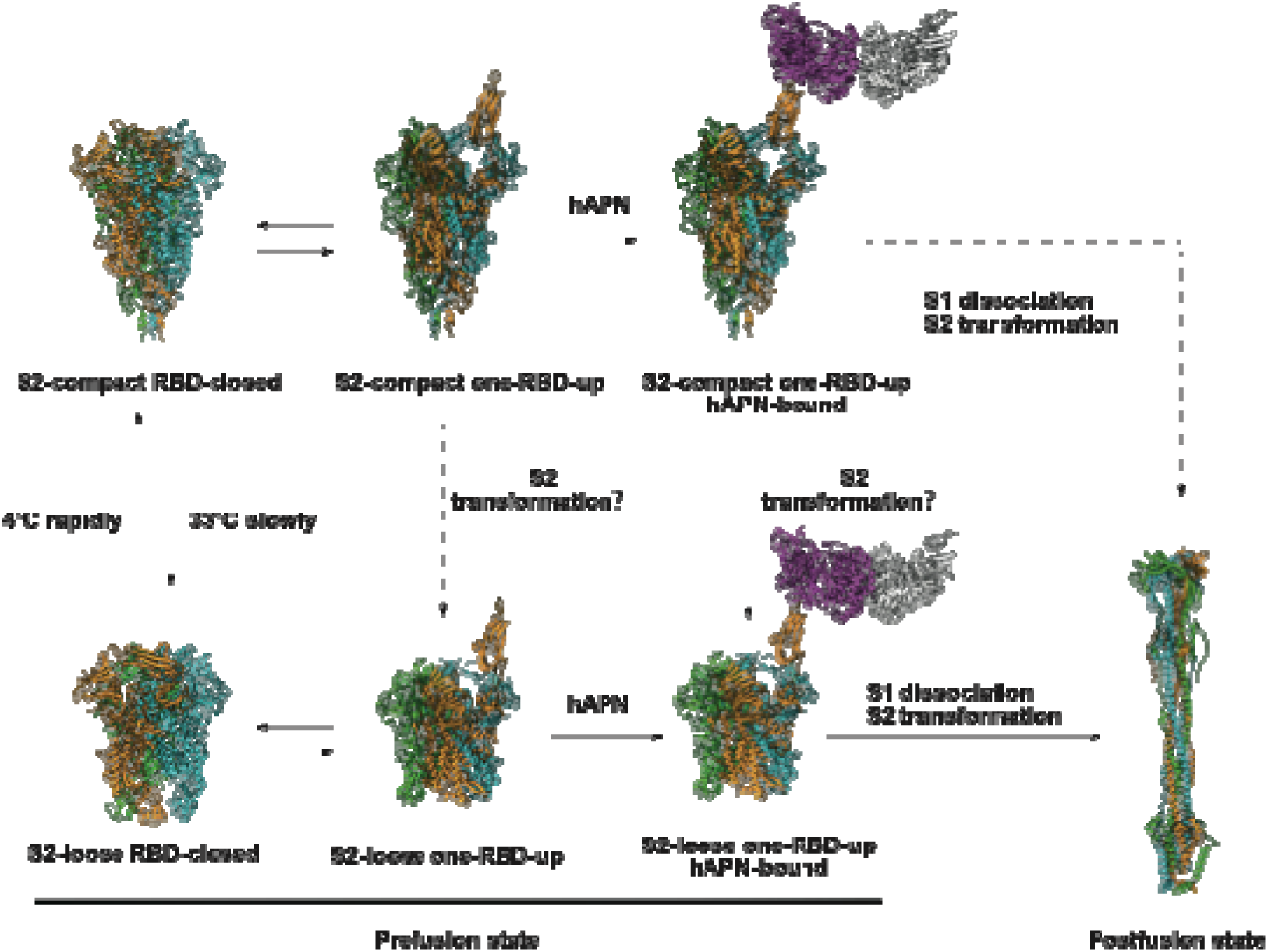
Model for temperature- and receptor-dependent activation of the HCoV-229E. **S.** Proposed model illustrating the coordinated roles of temperature and receptor engagement in regulating conformational transitions of the HCoV-229E S on-virions. At low temperature (4lJ), prefusion S trimers predominantly adopt an RBD-closed, S2-compact conformation. Exposure to physiological temperature (33lJ) gradually shifts the equilibrium toward the S2-loose conformation. In both S2 conformations, S can sample RBD-up states, enabling engagement with hAPN and formation of Solo or Gemini S-hAPN complexes. Receptor binding promotes S1 dissociation and S2 rearrangements, driving membrane fusion and conversion to the postfusion state. This model highlights the synergistic effects of temperature, receptor interaction, and incubation time in governing alphacoronavirus spike activation and viral entry. The RBD-closed, S2-compact model was determined from the on-virion prefusion S of HCoV-229E_live_-hAPN-33lJ_8h, while the RBD-closed, S2-loose model was from recombinant 2P S model (PDB: 6U7H) and the two S2-loose, one-RBD-up models (hAPN-unbound or hAPN-bound) were from recombinant S2-loose, S-hAPN model (PDB: 8WDE). The two S2-compact, one-RBD-up models (hAPN-unbound or hAPN-bound) were built by superimposing the recombinant apo hAPN ectodomain model and the up-RBD component of recombinant S2-loose, S-hAPN model (PDB: 8WDE) onto on-virion RBD-closed, S2-compact model with some modifications. The postfusion S model was from the homologous SARS-CoV-2 structure (PDB: 8FDW). The HCoV-229E S-trimer models are depicted with three protomers colored cyan, green, and orange. The hAPN monomers are colored magenta when bound to S, and white when unbound.

The temperature-dependent conformational behavior of HCoV-229E S is particularly relevant to viral transmission and infection in the human upper respiratory tract during winter. In cold environmental conditions, virions predominantly harbor S2-compact S, which may help preserve the integrity of prefusion S. Upon inhalation, exposure to warmer temperatures in the upper respiratory tract promotes a gradual transition toward the S2-loose conformation. In the presence of hAPN, both S2-compact and S2-loose S with exposed RBDs may engage receptors and initiate entry. Our observation that prolonged cultivation of HCoV-229E at physiological temperature yields virions enriched in S2-loose S is consistent with the typical 2-5 days incubation period of infection^9^ and suggests that accumulation of S2-loose S may facilitate viral dissemination at later stages of infection. Conversely, release of progeny virions into cooler environments would favor rapid reversion to the S2-compact conformation, thereby preventing premature transition to the postfusion state.

Finally, our findings have implications for vaccine and antiviral design against alphacoronaviruses. The ability of coronavirus S to switch between RBD-up and RBD-closed states allows viruses to balance receptor engagement with immune evasion. In highly pathogenic betacoronaviruses such as SARS-CoV-2, a substantial fraction of prefusion S adopt RBD-up conformations^33^, likely contributing to enhanced infectivity and transmissibility^42^. In contrast, due to distinct S1 quaternary packing, only a small fraction of RBD-up S is present on apo HCoV-229E virions. Instead, physiological temperature promotes a slow shift toward an S2-loose conformation enriched in RBD-up states, potentially fine-tuning viral adaptability within the host. These differences may underlie the distinct immunogenic profiles of alpha- and betacoronaviruses. Whereas potent neutralizing antibodies (nAbs) against SARS-CoV and SARS-CoV-2 predominantly target the RBD^43^, nAbs elicited by HCoV-229E infection more frequently recognize the NTD^44,45^. Moreover, the pronounced conformational flexibility of the HCoV-229E S2 subunit suggests that S2 represents an additional antigenic target. Compared with the S2-compact conformation, the S2-loose conformation likely exposes otherwise occluded epitopes recognized by S2-directed antibodies^45^. Given that prolonged exposure to physiological temperature favors the S2-loose conformation, immunogens based on this conformation may represent a promising strategy for alphacoronavirus vaccine development.

## Materials and Methods

### Cells and viruses

The laboratory strain HCoV-229E (ATCC VR740) was obtained from the China Center for Type Culture Collection (CCTCC) and stored at −80 until further use. Virus propagation was carried out in Huh-7 cells (the Cell Resource Center of the Institute of Basic Medical Sciences, Beijing, China) cultured at 33 for 5 days. Cell culture supernatants were harvested and clarified by low-speed centrifugation (3,000 g, 30 minutes, 4) to remove cellular debris.

For cryo-ET, virions were isolated using a discontinuous sucrose cushion, following previously described protocols^46^. Briefly, clarified supernatants were layered onto a discontinuous sucrose gradient consisting of 30% and 50% (w/v) sucrose in phosphate-buffered saline (PBS, pH 7.4) and subjected to ultracentrifugation in a Beckman SW32.1 rotor at 100,000 g for 3 h at 4. The virus band at the sucrose interface was collected using BioComp Gradient Station 108 (BioComp) and concentratedan Amicon Ultra Centrifugal Filter (Millipore, 100 kDa cutoff) at 4.

For fixation prior to isolation, an appropriate volume of clarified supernatant was loaded into the dialysis bag and immersed in a 15-fold excess volume of 4% PFA (Beijing Solarbio Science & Technology Co., Ltd.). Then samples were placed on a magnetic stirrer and incubated overnight at either 4 or 33 to allow diffusion-driven fixation. Fixed samples were then subjected to purification as described above. For fixation after isolation, concentrated virions were mixed with an equal volume of 8% PFA to a final concentration of 4% and incubated for 1.5 h. All procedures involving virus propagation, purification, concentration, and inactivation were conducted in a biosafety level 2 (BSL-2) laboratory at Tsinghua University.

### Receptor expression and purification

Expression and purification of hAPN ectodomain were performed as described^20^. Plasmids encoding the hAPN ectodomain fused to a C-terminal Fc tag were transiently transfected into HEK293F cells using PEI MAX (Polysciences) at a plasmid-to-PEI mass ratio of 1∶3. Four days post-transfection, cell culture supernatants were harvested and centrifuged at 4,000 rpm for 20 min to remove cellular debris. The hAPN-Fc fusion protein was purified by Protein A affinity chromatography using a HiTrap Protein A HP column (Cytiva) and subsequently concentrated using an Amicon Ultra centrifugal filter unit (Millipore, 30 kDa cutoff) at 4.

### Cryo-ET sample preparation

Live virions were mixed with 10 nm BSA (bovine serum albumin)-gold tracer (Aurion) and 4 μL of the mixture was applied to glow-discharged Quantifoil R2/1 or R2/2 holey carbon copper grids. Grids were blotted for 3 s and plunge-frozen into liquid ethane using a Cryo-Plunger 3 (Gatan Inc.) in a BSL-2 laboratory at Tsinghua University. For hAPN-incubated samples, live virions were incubated with soluble hAPN under the following conditions prior to grid preparation: 0.22 mg/mL hAPN at 33 for 8 h; 0.28 mg/mL hAPN at 4 for 24 h; or 0.28 mg/mL hAPN at 33 for 24 h. Inactivated virions were similarly mixed with 10 nm BSA-gold tracer and 4 μL of the mixture was applied to glow-discharged Quantifoil R2/1 holey carbon copper grids (200 mesh) using a Vitrobot Mark IV (Thermo Fisher Scientific) operated at 100% humidity and 8. Grids were blotted for 4.5 s, plunge-frozen in liquid ethane, and stored in liquid nitrogen until imaging.

### Cryo-ET data acquisition

Cryo-ET data were collected on a Titan Krios transmission electron microscope (Thermo Fisher Scientific) operated at 300 kV and equipped with a K3 direct electron detector (Gatan Inc.) and a BioQuantum energy filter (20 eV slit width). Tilt series were recorded in super-resolution mode at nominal magnifications of 64,000× or 81,000×, corresponding to calibrated pixel sizes as indicated in Supplementary Materials. Tilt series were acquired using SerialEM^47^, with dose-symmetric or PACEtomo schemes from −60° to 60° at 3° steps, eight or ten frames per tilt angle, a total accumulated dose of 131.2 e /Å², and defocus values ranging from −2 to −4 μm.

### Cryo-ET data processing

Tilt series preprocessing was performed using the high-throughput FlyTomo pipeline^34^, including removal of dark frames and optimization of gold fiducial localization. Prefusion S particles were automatically identified and segmented using a previously described method ^34^. Particle coordinates were calculated from the output files and were converted into a Dynamo-readable format. Initial S orientations were estimated based on vectors normal to the local viral membrane.

To classify S2 conformations, we performed STA using Dynamo^48^ and Relion4^49^. Particles were extracted from 8 × binned tomograms into subtomograms (box size 48 × 48 × 48) in Dynamo. A low-resolution 8 × binned prefusion S map from no-orientation-refinement reconstruction of the initial-orientation particles was used as the initial template for their alignment applying C3 symmetry. After removing particles with tilt angles exceeding 90°, the coordinates of the remaining particles were converted to Relion4-format for further refinement. Classification on the overall S particles identified S2-compact and S2-loose conformations. For RBD conformation classification, a focused-classification was performed. A mask focusing on the RBD-related region was generated followed by the 3D-classification without alignment. The class of RBD-up-like density was selected and reconstructed into an RBD-up S. For HCoV-229E_live_-hAPN-33 _8h, manual distinction classified the RBD-up S into hAPN-bound S and hAPN-unbound S.

For S-hAPN conformation classification, 6,223 manual-picked S-hAPN particles from 370 tomograms on HCoV-229E_live_-hAPN-33 _24h were extracted from 8 × binned tomograms into subtomograms (box size 64 × 64 × 64) in Dynamo. A low-resolution 8 × binned prefusion S-hAPN map from no-orientation-refinement reconstruction of the initial-orientation particles was used as the initial template for their alignment applying C1 symmetry. Subsequently, the particles were subjected to multi-reference alignment imposing C1 symmetry using the Solo S-hAPN map and using artificial Gemini S-hAPN map as the templates (lowpassed to 50 Å resolution). The S-hAPN complexes were classified into solo S-hAPN complexes or Gemini S-hAPN complexes. For Gemini S-hAPN complexes, the coordinates were moved to the center of the complex with overlapped particles removed, followed by further alignment with C2 symmetry imposed.

For the postfusion S reconstruction, 1,750 manual-picked postfusion S particles from 370 tomograms on HCoV-229E_live_-hAPN-33 _24h were extracted from 8 × binned tomograms into subtomograms (box size 48 × 48 × 48). A low-resolution 8 × binned postfusion map from no-orientation-refinement reconstruction of the initial-orientation particles was used as the initial template for their alignment applying C3 symmetry in Dynamo. Then the particles were extracted from 4× binned tomograms into boxes of 96 × 96 × 96 voxels for further alignment in Dynamo.

### Single-particle cryo-EM of hAPN

3 μL purified hAPN ectodomain (0.3 mg/mL) was applied to glow-discharged holey carbon grids (Quantifoil, R1.2/1.3, 300 mesh). Grids were blotted using Vitrobot Mark IV (Thermo Fisher Scientific) with a 4 s blotting time, a force level of −2 at 100% humidity and 8 and were then immediately plunged into liquid ethane cooled by liquid nitrogen. Data were collected on a Talos Arctica electron microscope (Thermo Fisher Scientific) operated at 200 kV and equipped with a K2 Summit direct electron detector (Gatan Inc.) with a calibrated pixel size of 0.94 Å. Movies were dose-fractioned into 32 frames with a total dose of 50 e^-^_/_A^2^ on the samples. The defocus values used during data collection varied from −1.5 to −1.8 μm. All images were collected using the SerialEM automated data collection software package. Micrographs were motion-corrected and dose-weighted using MotionCor2^50^. Subsequently, non-dose-weighted sums of power spectra were used to estimate the CTF with CTFFIND4^51^.

Then 579 images were processed using cryoSPARC^52^. The particles were auto-picked using Template picker and extracted with a box size of 200 pixels. 146,162 auto-picked particles were screened and selected by reference-free 2D Classification, while the low -resolution and wrong classes were deleted. The remaining 89,513 particles were then used for ab-initio reconstruction. Then the class (72,577 particles) with more intact density was selected for final Non-uniform 3D Refinement with C3 symmetry. The final density map was reconstructed to an overall resolution of 3.75 Å, according to the gold-standard Fourier shell correlation (FSC) threshold of 0.143. The local resolution of the final density maps was analyzed and estimated by the Local Resolution Estimation tool in cryoSPARC.

### Model building and refinement

To interpret the S structure, the recombinant S models were docked into on-virion density map using Chimera and ChimeraX. For model building, the initial atomic models for on-virion RBD-closed, S2-compact S and hAPN ectodomain structures were predicted using cryoNET^53^. These models were then manually adjusted in Coot, with glycans built manually into recognizable glycosylation sites based on the density. Next, models were real-space refined in PHENIX, followed by iterative correction and rebuilding in Coot. The final models were validated using comprehensive validation tools implemented in PHENIX. All geometry statistics and validation details are recorded in Extended data Tables.

### Viral titer assays

The titer of HCoV-229E was determined by focus forming assay (FFA) as previously described^54^. Following virus adsorption at 33 for 1 h, unbound virus was removed by washing with PBS. At 2 dpi, foci were visualized using TrueBlue Peroxidase Substrate (Kirkegaard & Perry Lab Inc.), and counted with an A-08 vSpot Spectrum reader (AID). Viral titers were calculated and expressed as focus forming unit (FFU)/mL.

For Fig. 1a, virus stocks were propagated under two conditions (at 33 or 37 for 5 days). This paired infection experiment was independently repeated four times. For each group, the viral titer was finally determined in triplicate using eight wells (dilution factor 10^3^) of a 96-well plate per replicate, and the average value was calculated. FFA results were analyzed and plotted using GraphPad Prism. Paired t-test revealed a statistically significant difference between the two propagation conditions (** indicates *P* value = 0.0070 < 0.01).

For Fig. 3c, Huh-7 cells were infected with virus stocks that had been pre-incubated under two conditions (maintained at 33 or 4 for 12 h). This paired temperature-treatment experiment was independently repeated four times. The virus stocks used for the 33 condition were the same as those in Fig. 1a. For each group, the viral titer was finally determined in triplicate using eight wells (dilution factor 10^3^) of a 96-well plate per replicate, and the average value was calculated. FFA results were analyzed and plotted using GraphPad Prism. Paired t test showed differences between two conditions were statistically significant (** indicates *P* value = 0.0048 < 0.01).

### Quantification and statistical analysis

For Figs. 1b, 2f, 3a,b, and 4c, the statistics are recorded in Extended Data Fig. 2, Extended Data Tables 1,2. For Fig. 4a, quantification of manually picked S-hAPN complexes relative to total prefusion S and virion counts revealed the following ratios (S-hAPN particles/total prefusion S/virion count): 143/17,658/1,183 for HCoV-229E_live_-hAPN-33 _8h; 461/5,055/611 for HCoV-229E_live_-hAPN-4 _24h; and 6,223/26,385/4,101 for HCoV-229E_live_-hAPN-33 _24h. The total prefusion S particles were picked using the FlyTomo pipeline. For Fig. 4d, 3,568 particles of solo S-hAPN complexes and 2,026 particles of Gemini S-hAPN complexes were used for statistic. Each Gemini S-hAPN complex contains two S trimers. For Fig. 4e, statistical analysis included 4,784 prefusion S and 157 postfusion S from 340 HCoV-229E_fixed_-4 _24h virions, 4,081 prefusion S and 121 postfusion S from 313 HCoV-229E_fixed_-33 _24h virions, 5,055 prefusion S and 398 postfusion S from 585 HCoV-229E_live_-hAPN-4 _24h virions, and 1,942 prefusion S and 505 postfusion S from 270 HCoV-229E_live_-hAPN-33 _24h virions. In apo virions, the average numbers of prefusion S were ∼14 ± 9 (mean ± SD, HCoV-229E_fixed_-4 _24h virions) and ∼13 ± 9 (HCoV-229E_fixed_-33 _24h virions), while the postfusion S numbers were both ∼0 ± 1. In hAPN-incubated virions, the average prefusion S numbers were ∼9 ± 6 (HCoV-229E_live_-hAPN-4 _24h) and ∼7 ± 5 (HCoV-229E_live_-hAPN-33 _24h), with corresponding postfusion S numbers of ∼1 ± 1 and ∼2 ± 2, respectively.

For Extended Data Fig. 3a, the diameters of virions were estimated using 876, 852, and 1,175 ellipsoidal models for the HCoV-229E_live_-4 _8h, HCoV-229E_live_-33 _8h, and HCoV-229E_live_-hAPN-33 _8h samples, respectively. For Extended Data Fig. 3b, the proportion of S2-compact prefusion S on intact virions was analyzed using 5,000 S from 450 virions (HCoV-229E_live_-4 _8h), 10,562 S from 766 virions (HCoV-229E_live_-33 _8h), and 17,582 S from 1,169 virions (HCoV-229E_live_-hAPN-33 _8h). For Extended Data Fig. 3c, the tilt angles of prefusion S were calculated as previously described for SARS-CoV-2^33^. Statistical analysis included 4,368 S2-compact S (within 90 degrees) from intact HCoV-229E_live_-4 _8h virions; 3,794 S2-compact and 1,130 S2-loose S from HCoV-229E_live_-33 _8h virions; and 4,473 S2-compact and 2,032 S2-loose S from HCoV-229E_live_-hAPN-33 _8h virions.

## Acknowledgments

We thank Dr. Jianlin Lei, Dr. Fan Yang and Dr. Niyun Zhou from the cryo-EM Facility, Technology Center for Protein Sciences, Tsinghua University, for their support on cryo-EM data collection. We also thank the computational facility support on the cluster of Bio-Computing Platform and the BSL-2 public facility support (Tsinghua University Branch of China National Center for Protein Sciences Beijing).

## Declarations

### Author contributions

S.L. conceived and supervised the project. J.L. propagated the viruses, performed biochemical analysis and prepared the cryo-samples. J.L., C.P. and Z.Z. collected and processed the EM data. C.P., J.L., Z.Z., Y.S. and J.Y. analyzed the structures. J.L., Y.S., C.P., X.W. and S.L. wrote the manuscript. All authors critically revised the manuscript.

### Date and materials availability

All data needed to evaluate the conclusions in the paper are present in the paper or the Supplementary Materials. The data for this study have been deposited in the database Electron Microscopy Data Bank (EMDB).

