## Supplementary Info for "On-virion structural dynamics reveal temperature- and receptor-coordinated activation of an alphacoronavirus spike"

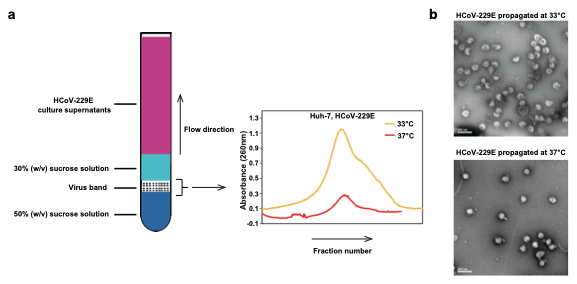
**Extended Data Fig. 1 | Characterization of propagation temperatures for HCoV-229E virions. a**, The absorption peaks of isolated HCoV-229E viral samples. The blue line represents the viral sample cultured at 33℃, while the orange line represents an equal volume of the viral sample cultured at 37℃. **b**, Negative staining images of the isolated HCoV-229E virions propagated at 33℃ and 37℃ from equivalent amounts of cell culture supernatants.

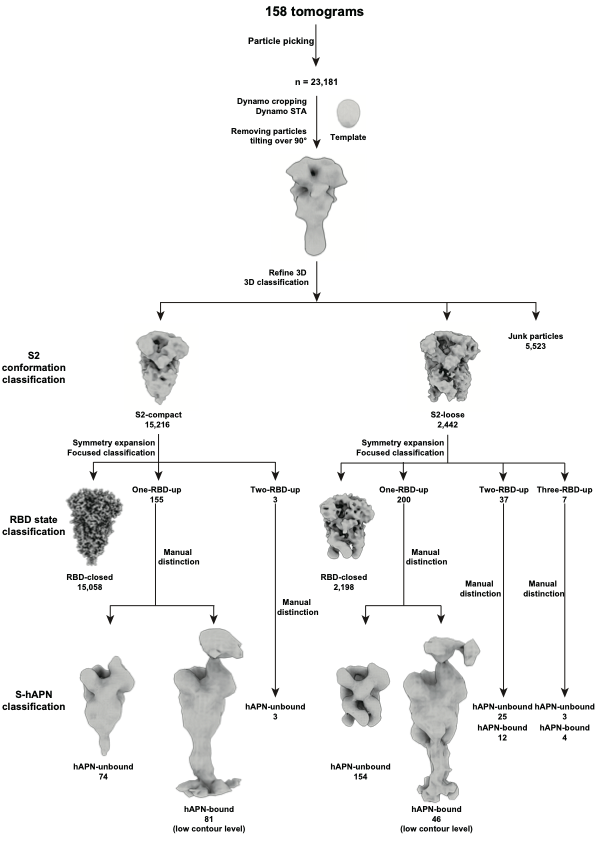
**Extended Data Fig. 2 | STA workflow for S on HCoV-229E virions.** Overview of the subtomogram averaging (STA) workflow used to analyze spike (S) trimers on HCoV-229E virions. The HCoV-229E_live_-hAPN-33℃_8h dataset is shown as a representative example. The same particle picking, alignment, and classification procedures were applied to all other datasets, including HCoV-229E_fixed_-33℃, HCoV-229E_fixed_-4℃_12h, HCoV-229E_live_-4℃_8h, HCoV-229E_live_-33℃_8h, HCoV-229E_fixed_-33℃_24h, and HCoV-229E_fixed_-4℃_24h.

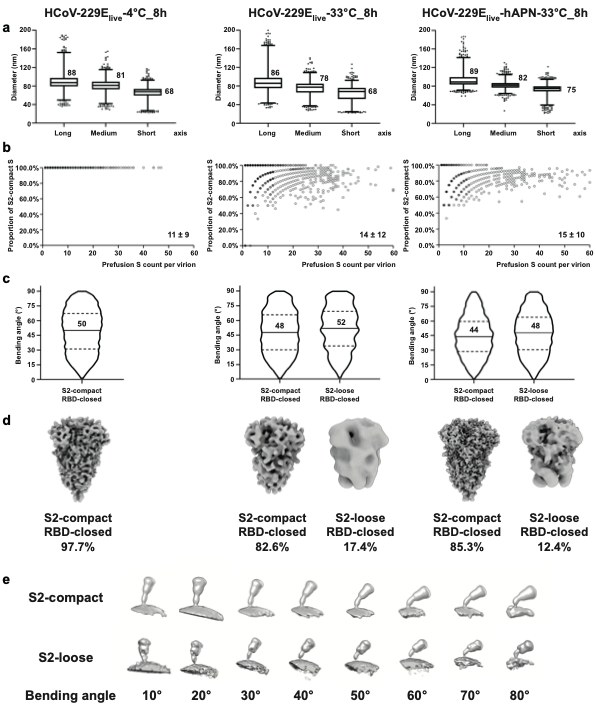
**Extended Data Fig. 3 | Statistics of S structures on apo or hAPN-incubated HCoV-229E virions. a**, Statistics of the dimensions of HCoV-229E viral envelopes. Boxplots display the 5% outliers, minimums, first quartiles, medians, third quartiles, and maximums of the data. The median diameters for the short, median, and long axis of the envelope are annotated. **b**, Distribution of prefusion S counts per virion and the corresponding proportions of S2-compact conformations. The coordinates of the scatter points correspond to the total number of prefusion S on their respective virions and the ratios of S2-compact S. The gray levels map to the point density. The average S counts for three samples are annotated (mean ± SD). **c**, Distribution of the prefusion S tilt angles on HCoV-229E virions reveals tilt ranges relative to the normal axis of the envelope. Violin plots indicate the first quartile, median, and third quartile. The median values are annotated. **d**, STA maps of RBD-closed prefusion S and their relative proportions across samples. **e**, STA maps of S in two S2 conformations on HCoV-229E_live_-hAPN-33℃_8h virions, displayed at multiple tilt angles.

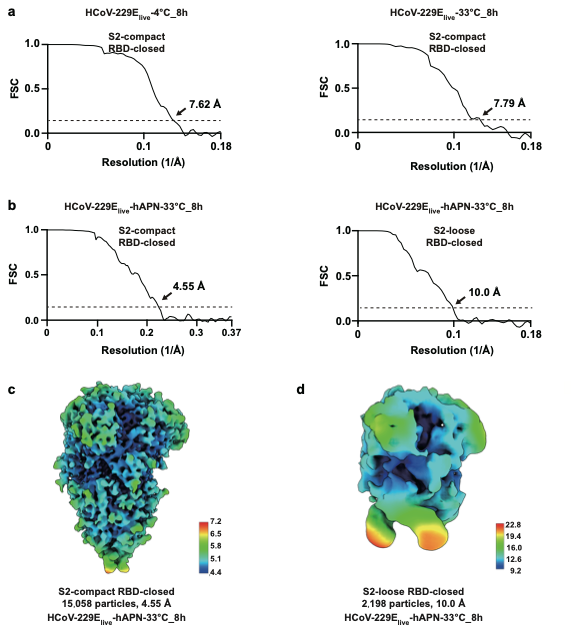

**Extended Data Fig. 4 | Local resolution of the RBD-closed prefusion S structure. a**, Resolutions of RBD-closed, S2-compact S structures from HCoV-229E_live_-4℃_8h and HCoV-229E_live_-33℃_8h samples were estimated from FSC curves, using a criterion of 0.143. **b**, Resolutions of RBD-closed, S2-compact and S2-loose, RBD-closed S structures from HCoV-229E_live_-hAPN-33℃_8h sample were estimated from FSC curves, using a criterion of 0.143. **c**, Map of the on-virion RBD-closed, S2-compact S from HCoV-229E_live_-hAPN-33℃_8h sample colored by its local resolution. **d**, Map of the on-virion S2-loose, RBD-closed S from HCoV-229E_live_-hAPN-33℃_8h sample. The map was colored by its local resolution.

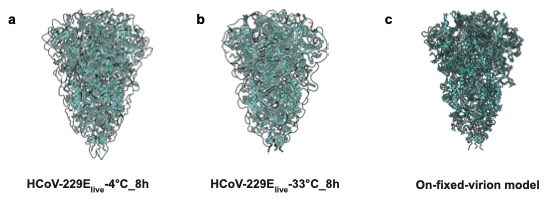

**Extended Data Fig. 5 | Structural comparison of on-virion RBD-closed, S2-compact S. a**, Rigid-body fitting of the RBD-closed, S2-compact S model derived from HCoV-229E_live_-hAPN-33℃_8h into the corresponding map from HCoV-229E_live_-4℃_8h virions. **b**, Rigid-body fitting of the same model into the RBD-closed, S2-compact map from HCoV-229E_live_-33℃_8h virions. **c**, Superposition of the RBD-closed, S2-compact S structure determined from HCoV-229E_live_-hAPN-33℃_8h virions and the structure resolved from 4℃-fixed apo virions (PDB: 9X6Z), with the RMSD = 1.097 Å.

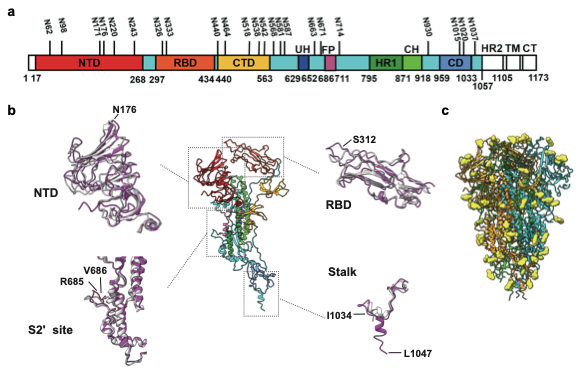
**Extended Data Fig. 6 | Structure of RBD-closed, S2-compact S on HCoV-229E virions. a**, Sequence schematic of HCoV-229E S, showing N-linked glycosylation sites. NTD, N-terminal domain; RBD, receptor binding domain; CTD, C-terminal domain; UH, upstream helix; S2' site, S2' cleavage site; FP, fusion peptide; HR1, heptad repeat 1; CH, central helix; CD, connector domain; HR2, heptad repeat 2; TM, transmembrane domain; CT, cytoplasmic tail. **b**, Structural features of the on-virion RBD-closed, S2-compact S. The structure was determined to 4.55 Å resolution from the HCoV-229E_live_-hAPN-33℃_8h virion by STA. Domains are colored as in **a**. Boxed regions including the RBD, NTD, S2', and stalk of on-virion model (magenta) are compared to those of the recombinant S model (light gray, PDB: 7CYC). In our on-virion structure, residue 176 adopts an additional glycan on NTD, the loops of RBM and S2' site show subtle changes; the stalk is determined until residue L1047. **c**, Glycosylation profile of the on-virion RBD-closed, S2-compact S, with N-linked glycans from one protomer shown in yellow.

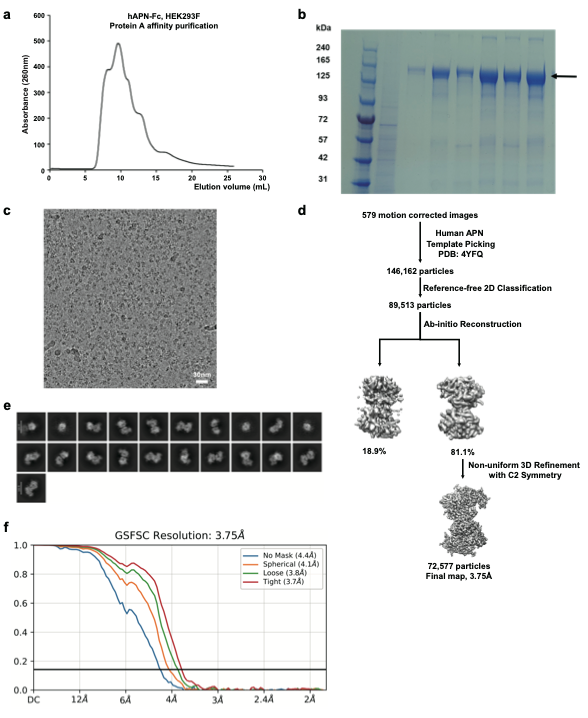
**Extended Data Fig. 7 | Purification, cryo-EM analysis of the hAPN ectodomain. a**, Protein A affinity chromatography of the hAPN ectodomain fused to a C-terminal Fc tag. **b**, SDS-PAGE analysis of the fractions from A. The black arrow indicates the position of the hAPN monomer. **c**, An exemplary cryo-EM raw micrograph of the hAPN dimers. Scale bar = 30 nm. **d**, Cryo-EM data processing workflow for hAPN. **e**, Representative 2D class averages of hAPN dimers. Scale bar = 100 Å. **f**, FSC curve used to estimate the resolution of the hAPN dimer reconstruction at the 0.143 criterion.

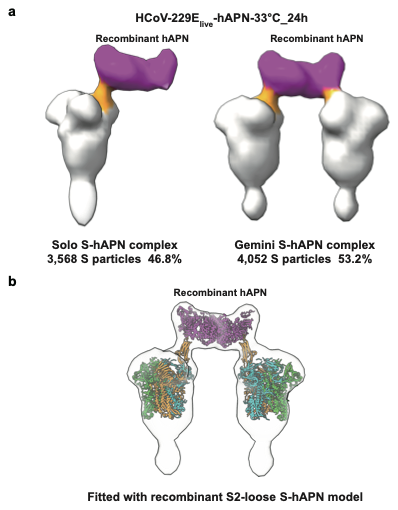
**Extended Data Fig. 8 | Structural fitting of on-virion S-hAPN complex on HCoV-229E_live_-hAPN-33℃_24h. a**, STA maps of on-virion S-hAPN complexes resolved from HCoV-229Elive-hAPN-33℃_24h virions. Left: solo S-hAPN; right: Gemini S-hAPN. RBDs (orange, up state) and hAPN dimers (magenta) are shown. **b**, Rigid-body fitting of the recombinant S2-loose S-hAPN complex model (PDB: 8WDE) into the on-virion Gemini S-hAPN density. The HCoV-229E S-trimer models are depicted with its three protomers colored cyan, green, and orange (RBD-up). The hAPN monomers are colored magenta (bound to S).

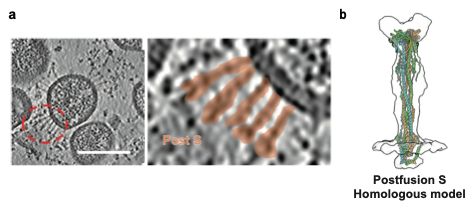

**Extended Data Fig. 9 | Postfusion S on HCoV-229E virions. a**, An exemplary tomogram slice (14 Å thickness) from HCoV-229E_live_-hAPN-33℃_24h virions showing postfusion S (red circle). Scale bar: 100 nm. **b**, STA map of the postfusion S resolved from HCoV-229E_live_-hAPN-33℃_24h virions, with rigid-body fitting of the homologous full-length SARS-CoV-2 postfusion S structure (PDB: 8FDW).

**Extended Data Table 1 | Cryo-ET data acquisition and reconstruction statistics of HCoV-229E**

| **Data acquisition** | | | | | | | | | |
| --- | --- | --- | --- | --- | --- | --- | --- | --- | --- |
| Microscope | Titan Krios | | | | | | | | |
| Magnification | 64,000/81000 | | | | | | | | |
| Voltage (kV) | 300 | | | | | | | | |
| Detector | Gatan K3 | | | | | | | | |
| Energy filter (eV) | 20 | | | | | | | | |
| Tilt schemes | Dose-symmetric/PACEtomo, dose-symmetric | | | | | | | | |
| Exposure  (e^-^/Å^2^) | 131.2 | | | | | | | | |
| Defocus range (μm) | -2.0 to -4.0 | | | | | | | | |
| Software | SerialEM | | | | | | | | |
| **Reconstruction** | | | | | | | | | |
| Software | Dynamo 1.1.532, Relion 4.0 | | | | | | | | |
| Samples | HCoV-229E_live_-4℃_8h | | HCoV-229E_live_-33℃_8h | | HCoV-229E_fixed_-4℃_12h | HCoV-229E_fixed_-33℃ | HCoV-229E_fixed_-4℃_24h | HCoV-229E_fixed_-33℃_24h | |
| Pixel size (Å)  (super-resolution) | 0.685 | | | | 0.68 | 0.54 | | | |
| No. of tomograms | 57 | | 67 | | 39 | 1,163 | 260 | 54 | |
| Structure datasets | S2-compact | | S2-compact | S2-loose | S2-compact | S2-loose | S2-compact | S2-compact | S2-loose |
|  | RBD-closed | RBD-up |  |  |  |  |  |  |  |
| Final no. of particles | 9,105 | 206 | 8,830 | 1,863 | 5,860 | 23,860 | 16,420 | 2,832 | 1,103 |
| Symmetry imposed | C3 | C1 | C3 | C3 | C3 | C3 | C3 | C3 | C3 |
| Final pixel size (Å) | 2.74 | 2.74 | 2.74 | 2.74 | 5.44 | 2.15 | 2.15 | 4.30 | 4.30 |
| Final resolution (Å) | 7.62 | N/A | 7.79 | N/A | N/A | 9.37 | N/A | N/A | N/A |
| Gold-standard | yes | no | yes | yes | no | yes | no | no | no |
| FSC threshold | 0.143 | N/A | 0.143 | N/A | N/A | 0.143 | N/A | N/A | N/A |
| *B* factor (Å^2^) | -348 | N/A | -372 | N/A | N/A | -160 | N/A | N/A | N/A |

**Extended Data Table 2 | Cryo-ET data acquisition and reconstruction statistics of hAPN-incubated HCoV-229E**

| **Data acquisition** | | | | | | | | | | | | |
| --- | --- | --- | --- | --- | --- | --- | --- | --- | --- | --- | --- | --- |
| Microscope | | | | Titan Krios | | | | | | | | |
| Magnification | | | | 64,000 | | | | | | | | |
| Voltage (kV) | | | | 300 | | | | | | | | |
| Detector | | | | Gatan K3 | | | | | | | | |
| Energy filter (eV) | | | | 20 | | | | | | | | |
| Tilt schemes | | | | Dose-symmetric/PACEtomo, dose-symmetric | | | | | | | | |
| Exposure (e^-^/Å^2^) | | | | 131.2 | | | | | | | | |
| Defocus range (μm) | | | | -2.0 to -4.0 | | | | | | | | |
| Software | | | | SerialEM | | | | | | | | |
| **Reconstruction** | | | | | | | | | | | | |
| Software | Dynamo 1.1.333, Dynamo 1.1.532, Relion 4.0 | | | | | | | | | | | |
| Samples | HCoV-229E_live_-hAPN-33℃_8h | | | | | | | HCoV-229E_live_-hAPN-4℃_24h | | HCoV-229E_live_-hAPN-33℃_24h | | |
| Pixel size (Å) (super-resolution) | 0.685 | | | | | | | 0.68 | | 0.68 | | |
| No. of tomograms | 158 | | | | | | | 67 | | 370 | | |
| No. of virions | 1,183 | | | | | | | 611 | | 4,101 | | |
| Structure datasets | S2-compact | | | | S2-loose | | | S-hAPN | Postfusion | Solo S-hAPN | Gemini S-hAPN | Postfusion |
|  | RBD-closed | One-RBD-up hAPN-unbound | One-RBD-up hAPN-bound | | RBD-closed | One-RBD-up hAPN-unbound | One-RBD-up hAPN-bound |  |  |  |  |  |
| Final no. of particles | 15,058 | 81 | 74 | | 2,198 | 154 | 46 | 461 | 594 | 3,568 | 2,026 | 1,750 |
| Symmetry imposed | C3 | C1 | C1 | | C3 | C1 | C1 | N/A | N/A | C1 | C2 | C3 |
| Final pixel size (Å) | 1.37 | 2.74 | 2.74 | | 2.74 | 2.74 | 2.74 | N/A | N/A | 10.88 | 10.88 | 10.88 |
| Final Resolution (Å) | 4.55 | N/A | N/A | | 10.0 | N/A | N/A | N/A | N/A | N/A | N/A | N/A |
| Gold-standard | yes | N/A | N/A | | yes | N/A | N/A | N/A | N/A | N/A | N/A | N/A |
| FSC threshold | 0.143 | N/A | N/A | | 0.143 | N/A | N/A | N/A | N/A | N/A | N/A | N/A |

**Extended Data Table 3 | Refinement and validation statistics of the cryo-ET RBD-closed, S2-compact S structure**

| **Refinement** | |
| --- | --- |
| Initial model used | Model predicated by cryoNET |
| Model resolution (Å) | 4.55 |
| FSC threshold | 0.5 |
| Model composition |  |
| Non-hydrogen atoms | 24,129 |
| Protein residues | 3,012 |
| Glycan residues | 69 |
| *B* factors (Å^2^) |  |
| Protein | 128.12 |
| Ligand | 147.36 |
| R.m.s deviations |  |
| Bond lengths (Å) | 0.004 |
| Bond angels (°) | 0.949 |
| **Validation** | |
| MolProbity score | 2.10 |
| Model vs. Map CC (volume) | 0.64 |
| Clashscore | 17.28 |
| Poor rotamers (%) | 0 |
| Ramachandran plot | |
| Favored (%) | 93.59 |
| Allowed (%) | 6.41 |
| Disallowed (%) | 0 |

**Extended Data Table 4 | Cryo-EM data acquisition and reconstruction statistics of soluble hAPN**

| **Data acquisition** | |
| --- | --- |
| Microscope | Talos Arctica |
| Magnification | 45,000 |
| Voltage (kV) | 200 |
| Detector | Gatan K2 |
| Pixel size (Å) | 0.94 |
| Frames | 32 |
| Exposure (e-/Å2) | 50 |
| Defocus range (μm) | -1.5 to -1.8 |
| Software | SerialEM |
| **Reconstruction** | |
| Software | CryoSPARC |
| Initial number of particles | 146,162 |
| Final number of particles | 72,577 |
| Symmetry imposed | C2 |
| Final Resolution (Å) | 3.75 |
| Gold-standard | Yes |
| FSC threshold | 0.143 |
| B factor (Å2) | 194 |

**Extended Data Table 5 | Refinement and validation statistics of the of soluble hAPN structure**

| **Refinement** | |
| --- | --- |
| Initial model used | Model predicated by cryoNET |
| Model resolution (Å) | 3.75 |
| FSC threshold | 0.5 |
| Model composition |  |
| Non-hydrogen atoms | 14,840 |
| Protein residues | 1,804 |
| Glycan residues | 20 |
| *B* factors (Å^2^) |  |
| Protein | 116.90 |
| Ligand | 138.27 |
| R.m.s deviations |  |
| Bond lengths (Å) | 0.004 |
| Bond angels (°) | 1.046 |
| **Validation** | |
| MolProbity score | 2.08 |
| Model vs. Map CC (volume) | 0.77 |
| Clashscore | 11.38 |
| Poor rotamers (%) | 0 |
| Ramachandran plot | |
| Favored (%) | 91.22 |
| Allowed (%) | 8.78 |
| Disallowed (%) | 0 |
